# PluMu - a Mu-like bacteriophage infecting *Actinobacillus pleuropneumoniae*

**DOI:** 10.1101/2024.02.07.579279

**Authors:** L Julia Bartsch, Roberto Fernandez Crespo, Yunfei Wang, Michael A Skinner, Andrew N Rycroft, William Cooley, David J Everest, Yanwen Li, Janine T Bossé, Paul R Langford

**Author notes:** Contributed equally (L Julia Bartsch, Roberto Fernandez Crespo).

## Abstract

*Actinobacillus pleuropneumoniae* is the causative agent of pleuropneumonia, an economically important lung disease of pigs. In draft genomes of two Cypriot clinical *A. pleuropneumoniae* isolates (MIDG3457 and MIDG3459), we previously identified single genomic regions with homology to Mu-like bacteriophage and presented preliminary evidence of active phage. Here, updated Phastest genomic analysis identified two loci in both MIDG3457 and MIDG3459 that were predicted to encode proteins with high homology to, and whose organization was characteristic of, Mu-like phages. Phylogenetically, the closest matches were with *Mannheimia* Vb and *Glaesserella* SuMu phages. Phastest scored the loci as “complete”, indicating they produced active phage. PCR amplification of the Mu-like phage *c* and *tail* genes from DNase-treated polyethylene glycol 8000 (PEG)-precipitated supernatants of MIDG3457 and MIDG3459 (grown in either Brain Heart Infusion-NAD or Grace’s Insect Medium-NAD broth) indicated the presence of intact virions. The phages from MIDG3457 and MIDG3459 were named PluMu 3457-1, 3457-2, and PluMu 3459-1 and PluMu 3459-2, respectively. Transmission electron microscopy (TEM) of PEG-precipitated supernatants of broth-grown MIDG3459 identified virions with icosahedral heads and tails, consistent with other Mu-like phages. We conclude that MIDG3459 produces an active Mu-like phage.

## 1. Introduction

*Actinobacillus pleuropneumoniae* is a bacterial pathogen that causes pleuropneumonia in pigs and is responsible for substantial losses in the worldwide pig industry [1,2]. Based on surface carbohydrates, principally capsule, there are 19 known serovars [2], and the bacterium is further classified as biovar 1 or biovar 2 if they are NAD-dependent or NAD-independent, respectively. Some serovars are considered more virulent than others. For example, serovars 1, 5, 9/11 and 16 are considered of high virulence, as they express ApxI and ApxII toxins; those expressing ApxII and ApxIII toxins, such as serovar 8, are considered of lesser virulence; and those expressing only one Apx toxin are considered the least virulent [3]. Control of *A. pleuropneumoniae* is through good husbandry practices, e.g., reduction/elimination of stressors (such as poor ventilation and/or temperature), vaccination, and/or antibiotic use [1]. First-generation killed whole-cell (bacterin) vaccines are the most widely used, but do not prevent colonization or transmission; second-generation Apx-toxin recombinant or toxoid vaccines have been associated with safety concerns; and third-generation live attenuated vaccines are in development [4]. Mobile genetic elements facilitate the spread of antimicrobial resistance (AMR) and small plasmids and integrative conjugative elements carrying resistance genes have been reported in isolates of *A. pleuropneumoniae* [5,6]. However, to our knowledge, no active phage has been described in *A. pleuropneumoniae* although recent Phaster [7] and ProPhage Hunter [8] analyses identified possible phage in some isolates [9,10]. However, no data was presented demonstrating the production of viral particles. In a whole genome sequencing (WGS) study of *A. pleuropneumoniae* isolates from throughout the world, preliminary manual genome annotation and experimental analysis identified single Mu-like phage loci in two Cypriot isolates, MIDG3457 and MIDG3459 – (hereafter referred to as 3457 and 3459) [11]. Mu phage was originally described in enterics, such as *Escherichia coli* [12] and *Shigella* [13], but Mu-like phages have also been reported in *Glaesserella* (*Haemophilus*) *parasuis* [14], and *Mannheimia haemolytica* [15], which like *A. pleuropneumoniae*, are members of the *Pasteurellaceae* family. Mu-phages are double-stranded DNA phages of the Caudovirales (tailed bacteriophage) order, and the Myoviridae family. Here we have reannotated the Mu-like phage loci in MIDG3457 and MIDG3459 and show the production of a Mu-like phage (named PluMu 3457 and PluMu 3459, respectively), as adjudged by DNA-sequence analysis, molecular analyses, and transmission electron microscopy (TEM).

## 2. Materials and methods

### 2.1. *A. pleuropneumoniae* strains and growth conditions

Three porcine lung sample isolates of *A. pleuropneumoniae* were used: MIDG2331 (hereafter referred to as 2331), and 3457 and 3459 were isolated in Cyprus (2011) and are serovar 11 and 1, respectively, as determined by PCR [2]. 2331 was isolated in the UK (1995) and is serovar 8 [16]. Genome analyses indicated the potential presence (in 3457 and 3459) [11] or absence (in 2331) [16] of a Mu-like prophage. All bacterial strains were cultivated either on 1.5% agar plates containing Brain Heart Infusion (BHI) (BD® Bacto, UK) supplemented with 0.01% NAD, or in liquid broth. BHI-NAD agar plates were placed in a static incubator at 37°C and 5% CO2 overnight. Liquid culture was in either BHI-NAD broth or Grace’s Insect Medium (Gibco™ Fisher Scientific, UK) [17], supplemented with 0.01% NAD. For growth curves, overnight *A. pleuropneumoniae* agar plate cultures were resuspended in 10 ml broth and triplicate suspensions, in 50 ml Falcon™ centrifuge tubes, were placed in a shaking incubator (37°C, 180 rpm) for up to 4 h with OD_600_ measurements taken at hourly intervals. Mean OD_600_ values across triplicates for each isolate were calculated. All chemicals were obtained from Sigma (UK) unless otherwise stated.

### 2.2. Genomic comparisons

Originally, single Mu-like phages were identified in 3457 and 3459 by manual whole genome annotation [11]. Here, an up-to-date annotation was done using the phage prediction program

Phastest [18]. The genomes of 3457 (ENA accession number: ERS155414) and 3459 (ENA accession number: ERS155416) were submitted to Phastest and, in each case, two intact phages were predicted (see Results). These DNA sequences were extracted from Phastest outputs and used in subsequent analyses. The genome sequence of 2331 (Genbank accession number: LN908249.1), which does not contain a Mu-like phage, has been previously published [16]. Confirmation of the Phastest predicted prophages in 3457 and 3459 was done against a nucleotide database of Mu-like viruses (taxid:10677) and using tblastx [19]. Further comparative analyses on nucleotide and protein sequence levels were performed using blastn, tblastx, and EasyFig [20]. Comparison of PluMu was also made against the following genomes: *Burkholderia* phage BcepMu (AY539836.1), *Escherichia* phage Mu (AF083977.1), *Escherichia* phage D108 (NC_013594.1 and *Glaesserella parasuis* phage SuMu (JF832915.1), *Mannheimia* phage vB (KP137440.1), *Pseudomonas* phage D3112 (AY394005.1), *Rhodobacter* phage RcapMu (NC_016165.1), *Shigella* phage SfMu (NC_027382.1), and *Vibrio* phage Martha 12B12 (NC_021070.1). Phylogenetic trees were constructed by uploading fasta files of phage genomes to the NGphylogeny server (www.ngphylogeny.fr) [21] and choosing the PhyM analysis suite. Newick output files were visualised using the Phylogenetic tReE viSualisaTion (PRESTO) [22].

### 2.3. Precipitation of viral supernatants

Liquid (10 ml) bacterial cultures were grown in a shaking incubator followed by centrifugation at 4000 g for 10 min and supernatants were passed through 0.22 μm (Millex, UK) filters. Filtered supernatants were precipitated by the addition of sterile 20% w/v polyethylene glycol-8000 (PEG) in 2.5 M NaCl in a 1:5 ratio. Precipitated samples were stored at 4°C overnight, centrifuged at 4500 g for 40 min at 4°C, and pellets resuspended in 150 μl of 1x phosphate-buffered saline (PBS, pH 7.4, Gibco, UK). Precipitated samples were either kept at 4°C for up to a week or stored at -20°C before further analyses.

### 2.4. PluMu production

The presence of PluMu in bacterial cultures was shown by specific PCR amplification of phage DNA regions using bacterial or PluMu DNA extracted from liquid cultures or derived-filtered or filtered PEG-precipitated supernatants. The QIAamp DNA Mini and Fast Cycling kits (QIAGEN, UK) were used for DNA extraction and PCR amplification, respectively, according to the manufacturer’s instructions. Three primer pairs were synthesised by Eurofins (Germany) and are listed in Table 1. Primers Mu c For and Mu c Rev target a fragment of the *c* gene that encodes the late transcription activator protein C that controls the four "late" promoters, Pmom, Plys, PI and PP [23]. Tail-_For and Tail_Rev primers amplify a fragment of a Mu tail protein gene. SodC_For and SodC_Rev amplify a fragment of the *A. pleuropneumoniae sodC* (encodes Cu, Zn superoxide dismutase) and the adjacent *asd* (encoding aspartate semialdehyde dehydrogenase) gene [24], which served as a bacterial control. Initial optimisation experiments established that a multiplex PCR with all primers was not possible. Therefore, duplicate samples were subjected to a multiplex PCR targeting *c* and *tail* genes and a separate *sodC*-targeting PCR. The PCR cycling conditions were as follows: an initial denaturation step at 95°C for 15 min; 30 cycles of 94°C for 30 sec, 57°C for 90 sec, and 72°C for 90 sec; followed by a final step of 10 min at 72°C.

**Table 1.**
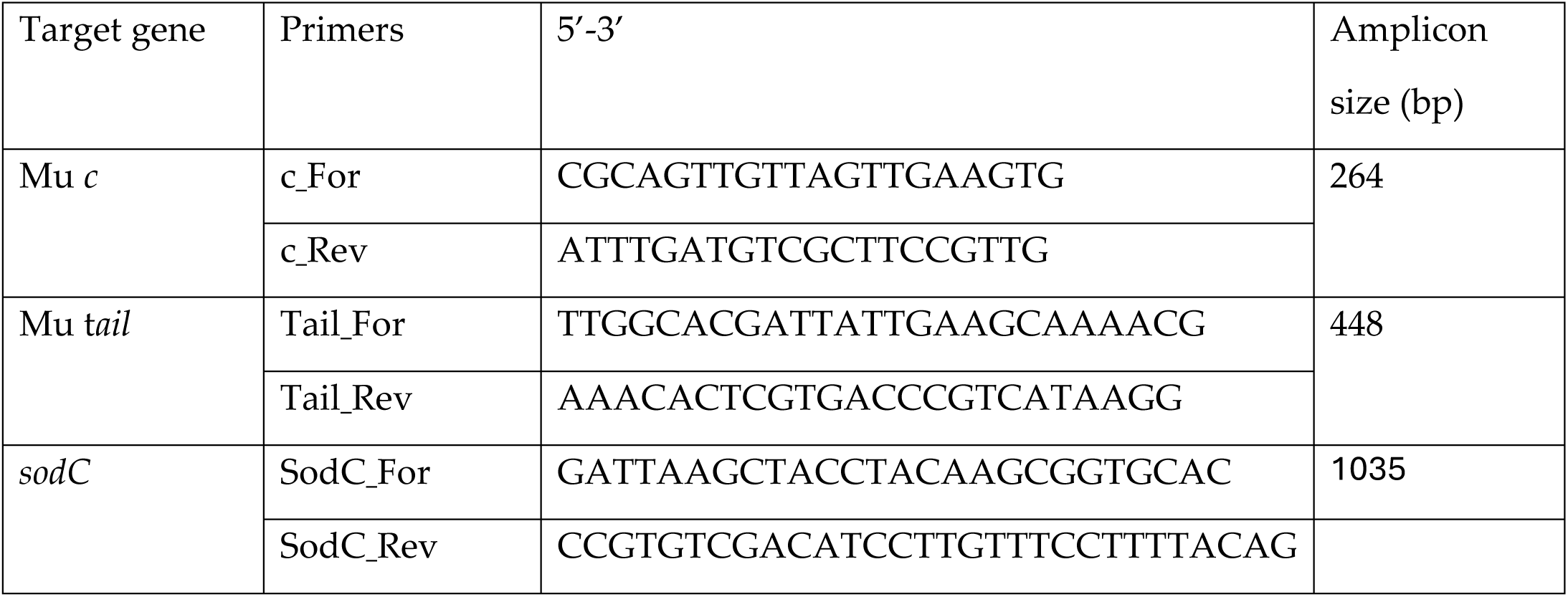
Sequences of primer pairs that were used in this study.

Amplicons were separated on 1% w/v agarose gels and stained using ethidium bromide (1 μg/ml). The GeneRuler 100 bp Plus DNA Ladder (Thermo Fisher Scientific, UK) was used as the marker. When required, gDNA derived from bacterial cultures, filtered supernatant, and filtered PEG-precipitated supernatants was removed using the TURBO DNA-free™ Kit (Invitrogen™ Ambion™, UK), according to the manufacturer’s instructions.

### 2.5. Transmission electron microscopy (TEM)

Filtered (0.22 μm) supernatants and PEG-precipitated supernatants of strains 2331, 3457, and 3459 were analysed. Samples (5 μl) were added to carbon-coated copper grids (Fisher Scientific, UK), stained with 2% phosphotungstic acid for 5 min, dried using Whatman filter paper and examined using a Tecnai bioTWIN transmission electron microscope at the Animal and Plant Health Agency core facilities.

## 3. Results

### 3.1. Genomic comparisons

A summary of the results of Phastest analysis on the previously identified manually annotated phage genome sequences from 3457 and 3459 is shown in Table 2. Phastest identified two DNA regions in the 3457 genome that encode proteins with high similarity to other Mu-like phages: Region 1 (PluMu 3457-1) of 41.9 kb and Region 2 (PluMu 3457-2) of 23 kb are predicted to encode 55 and 37 proteins, respectively. In the 3459 genome, Phastest also predicted two regions encoding proteins with high similarity to other Mu-like phages: Region 1 (PluMu 3459-1) of 36 kb and Region 2 (PluMu 3459-2) of 50 kb are predicted to encode 49 and 55 proteins, respectively. All four predicted phages were ranked by Phastest as “intact” (Table 1). An intact score is based on homology to known intact phage, the identity of the proteins (e.g., head, integrase, fiber, portal, terminase, protease or lysin), the size and number of proteins in the region, and the percentage of the phage-related and hypothetical proteins in the region. The maximum score is 150, and >90 is rated as intact. All but PluMu 3457-2 (score = 100), which has the smallest locus, have scores of 140 or 150.

**Table 2:**
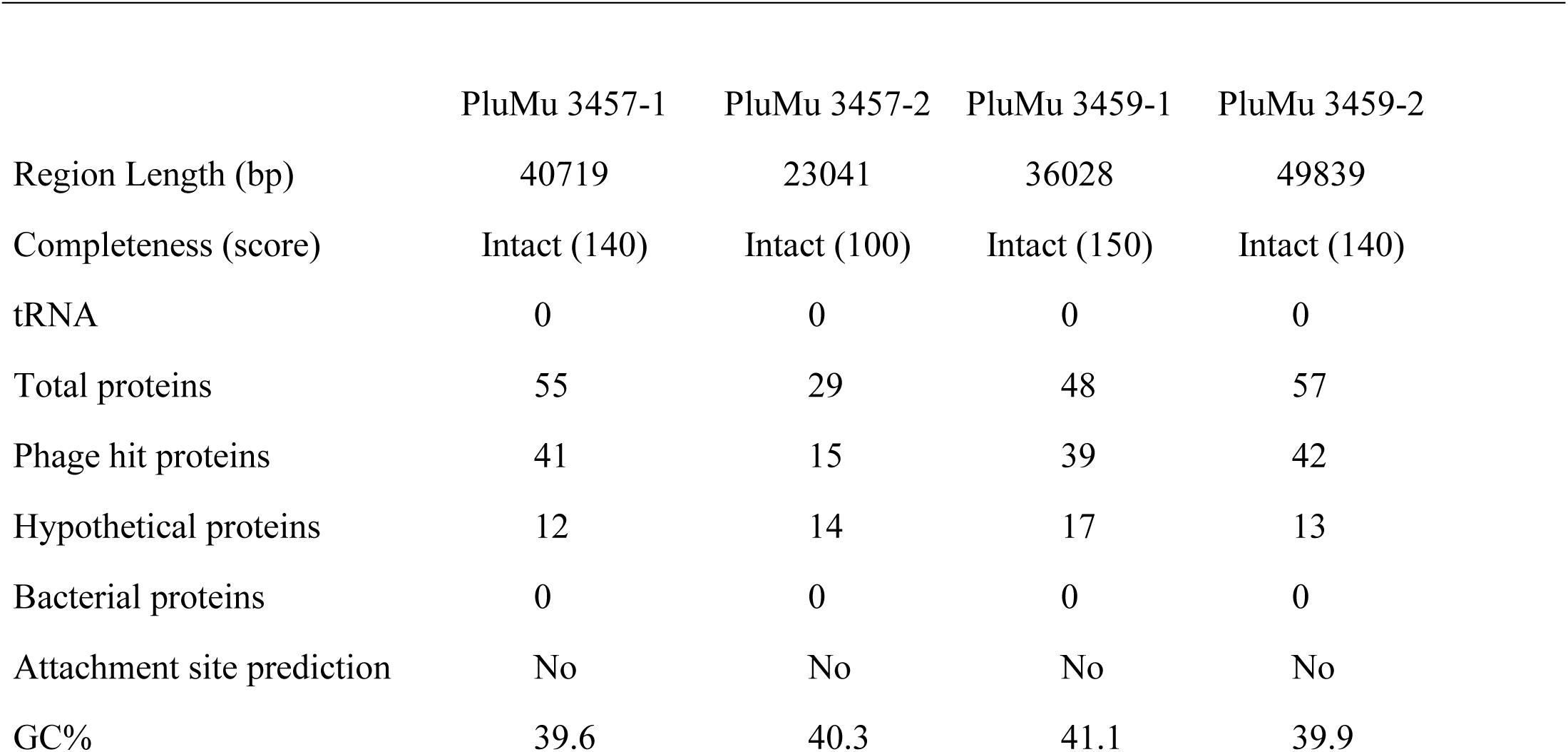
Summary features of PluMu phages identified the genomes of 3457 and 3459 by Phastest [18]. See text and Tables S1-4 for further details. A completeness score >90 = intact. The maximum score is 150.

|  | PluMu 3457-1 | PluMu 3457-2 | PluMu 3459-1 | PluMu 3459-2 |
| --- | --- | --- | --- | --- |
| Region Length (bp) | 40719 | 23041 | 36028 | 49839 |
| Completeness (score) | Intact (140) | Intact (100) | Intact (150) | Intact (140) |
| tRNA | 0 | 0 | 0 | 0 |
| Total proteins | 55 | 29 | 48 | 57 |
| Phage hit proteins | 41 | 15 | 39 | 42 |
| Hypothetical proteins | 12 | 14 | 17 | 13 |
| Bacterial proteins | 0 | 0 | 0 | 0 |
| Attachment site prediction | No | No | No | No |
| GC% | 39.6 | 40.3 | 41.1 | 39.9 |

The various PluMu phage sequences were inserted in different locations in the 3457 and 3459 genomes. For Mu-phage sequence orientations with X at the 5’ and Y at the 3’ end, the following insertion sites were found: PluMu 3457-1 is between *recO,* encoding a DNA repair protein and *qseB;* PluMu 3457-2 is between genes encoding the 50S ribosomal protein L25 and an uncharacterized protein; PluMu 3459-1 is between *rseA*, encoding a Sigma E negative regulatory protein [25], and an uncharacterized gene; PluMu 3459-2 interrupts the *thiH* gene, encoding the iminoacetate-synthetase, which is upstream of the gene encoding the 50S ribosomal protein L25. In each case, the most common blastp hits are against proteins encoded by Mu-like phages. The highest number of hits for PluMu 3457-1 (n=9) and PluMu 3459-2 (n=9) were against *Mannheimia* Vb (NC_028956), and PluMu 3457-2 (n=18) and PluMu 3459-1 (n=18) were against *Escherichia* D108 (NC_013594), Mu phages (Tables S1 and S2). Whilst there is comparatively little identity at the DNA level (Figure 1), there is high similarity at the protein level, especially at the 5’ end (see Figure 1), between the PluMu 3457-1 and PluMu 3459-1 (phages with the highest completeness score in 3457 and 3459, respectively), and *Mannheimia* Vb, *Escherichia* D108 and reference *Escherichia* (AF083977) Mu phages.

**Figure 1.**
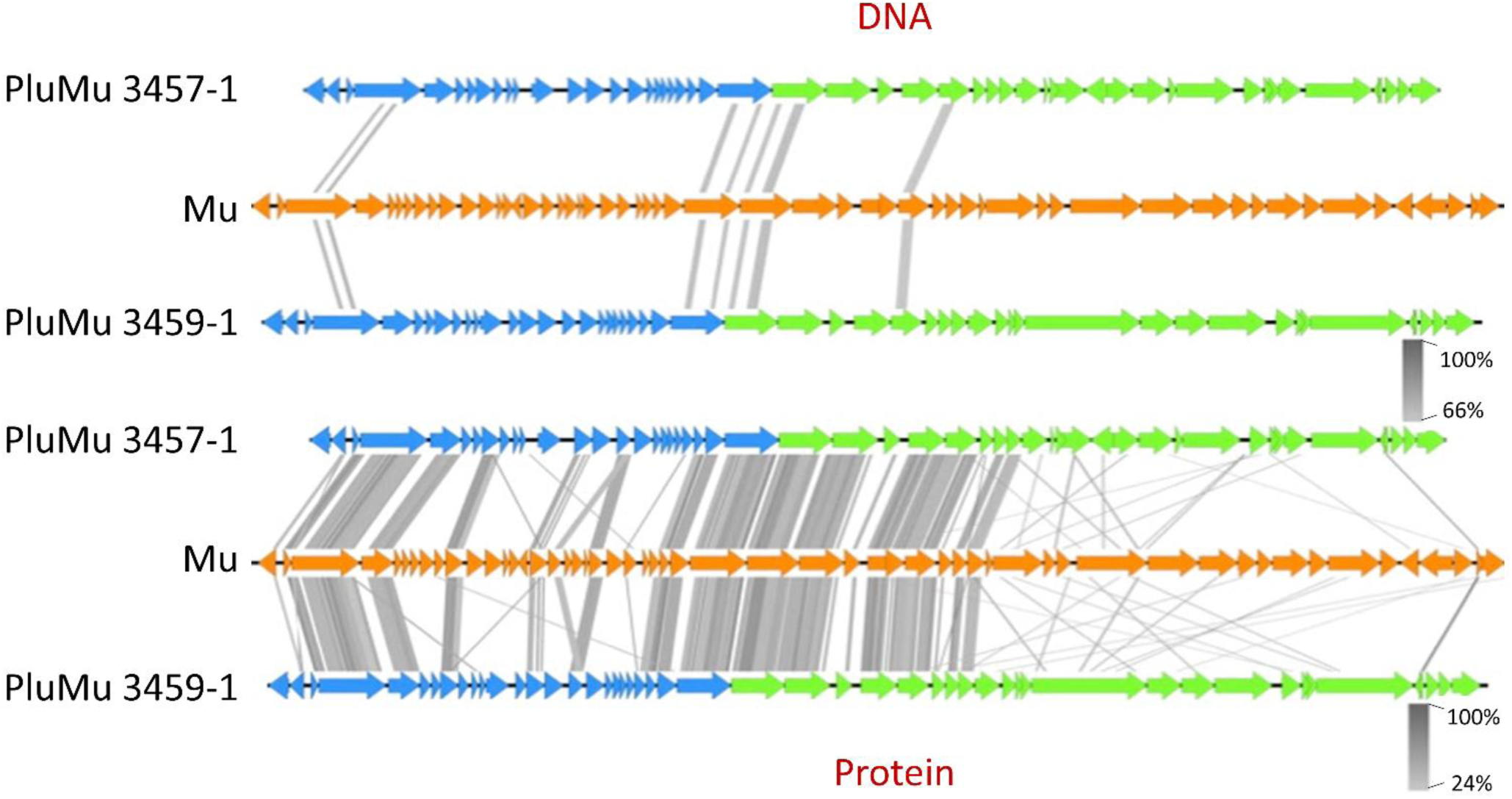
Sequence comparison of the reference *Escherichia* Mu bacteriophage (AF083977 and PluMu 3457-1 and 3459-1. Comparison is made at the nucleotide (top) and protein (bottom) levels. Genes in dark blue are those in the regulatory segment and those in green are related to viral particle synthesis. The grey scale on the bottom right represents the level of homology.

Phastest circular genomic output schematics of the organization of PluMu “intact” phage in 3457 and 3459 genomes are shown in Figure 2A-D, with details of the predicted CDSs, top blastp hits, and significant E-values in Tables S1-4. All have a typical Mu-like phage organization, i.e., latency, early, and middle (region finishing downstream of the *c* gene), followed by the late lysis-head-tail region, with key predicted genes and proteins as detailed by Harshey [26] and the ViralZone (https://viralzone.expasy.org/4356), available through the ViralZone server [27]. These include genes encoding integrases, repressors, anti-repressors, transposases, portal proteins, terminases, and those required for tail, and head formation.

**Figure 2.**
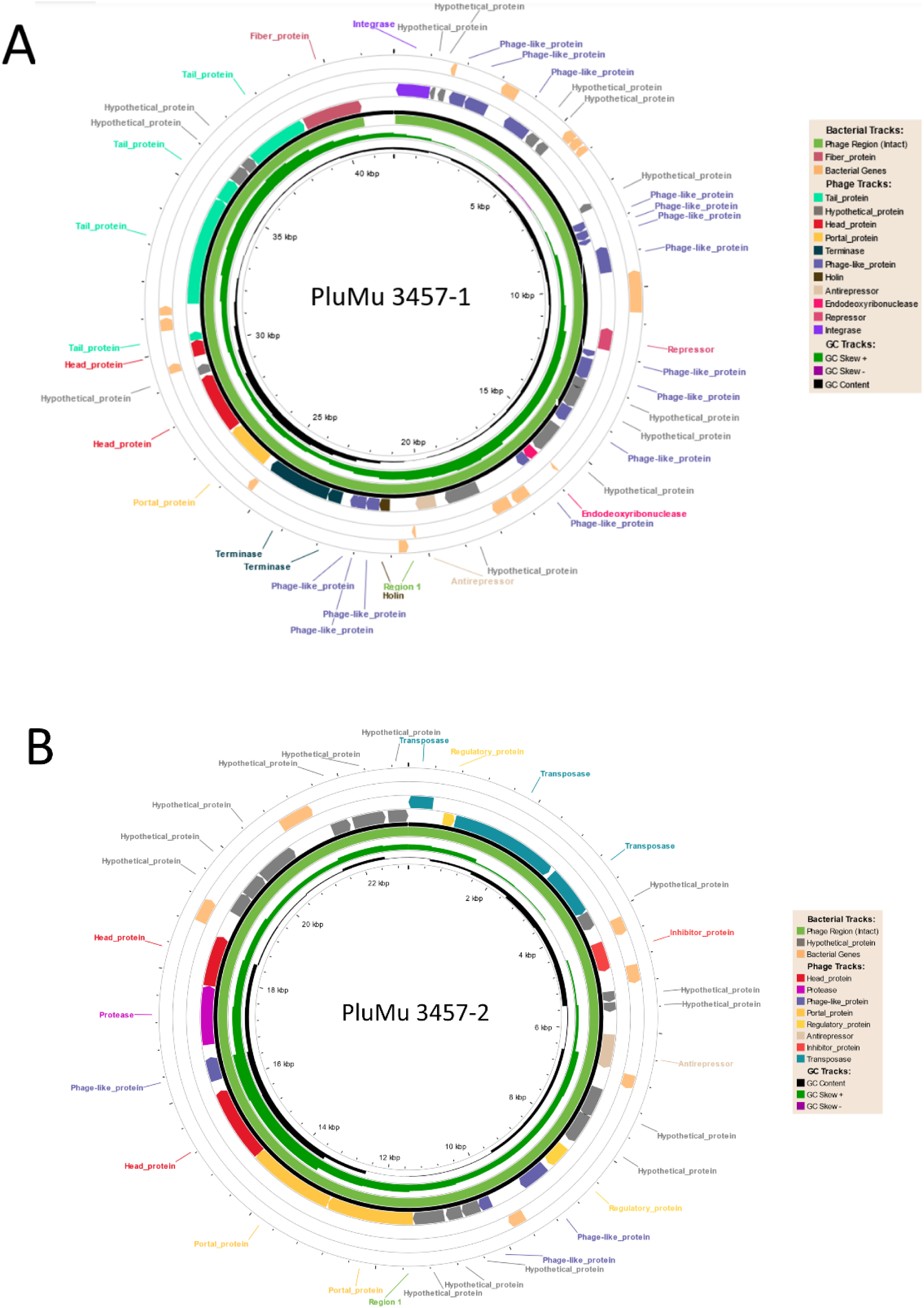
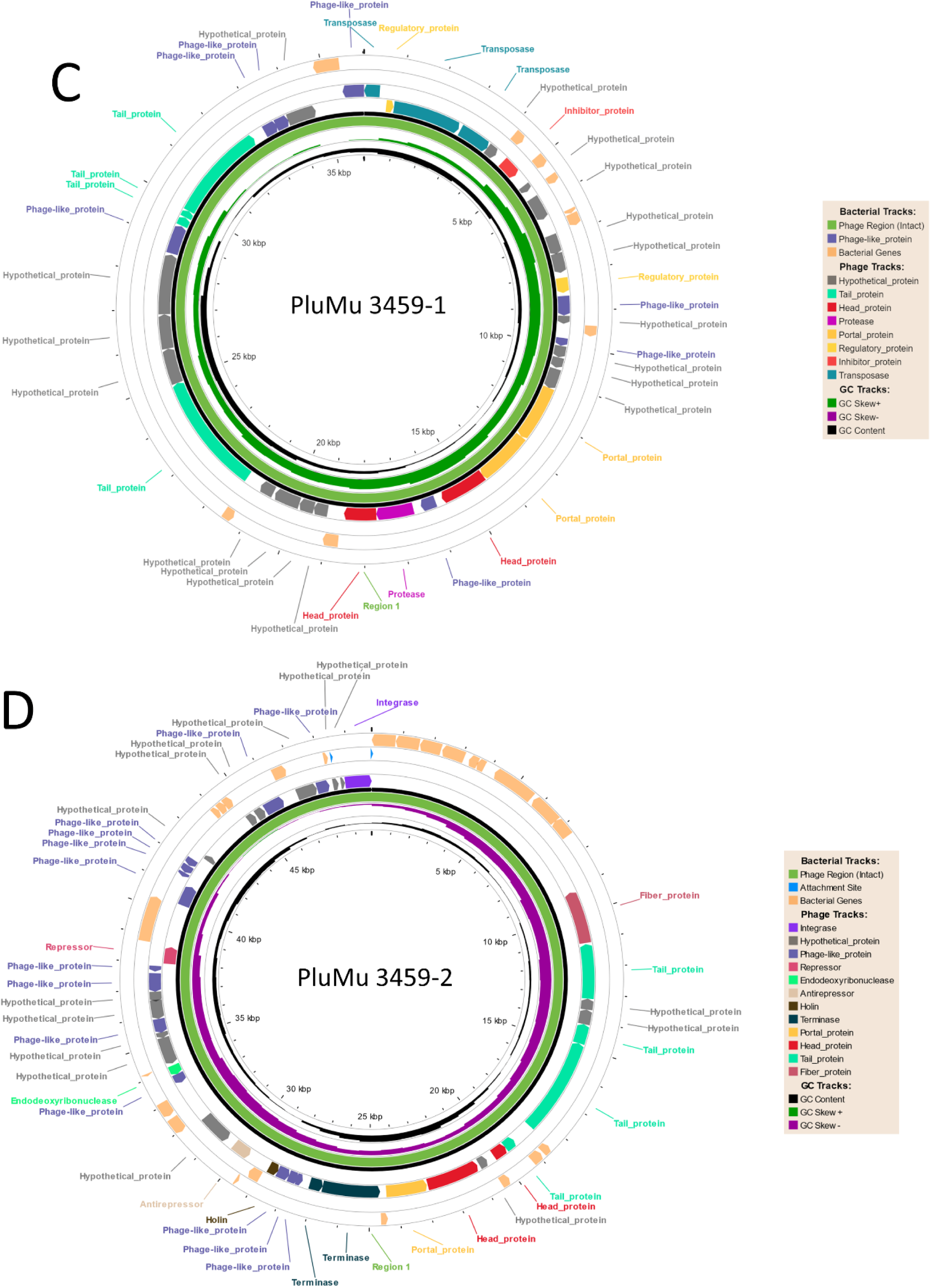
Phaster genome outputs showing organization and predicted functions of proteins of (A) PluMu 3457-1; (B) PluMu 3457-2; (C) PluMu 3459-1; and (D) PluMu 3459-2. See Tables S1-S4 for a list of predicted CDS, top blastp hits, and E-values, upon which the Figures are based.

A phylogenetic tree also demonstrated their relatedness to known Mu-phages (Figure 3). All four PluMu phages were closely related to each other, especially PluMu 3457-2 and PluMu 3459-1, and then to a subbranch including *Mannheimia* Vb and *Glaesserella* SuMu, the parent bacteria of which, like *A. pleuropneumoniae*, are members of the *Pasteurellaceae*. Thus, the Phastest genomic analyses indicated the presence of two phages each in 3457 and 3459, with characteristic Mu-like phage organization and intact completeness scores indicating their ability to produce active virions.

**Figure 3.**
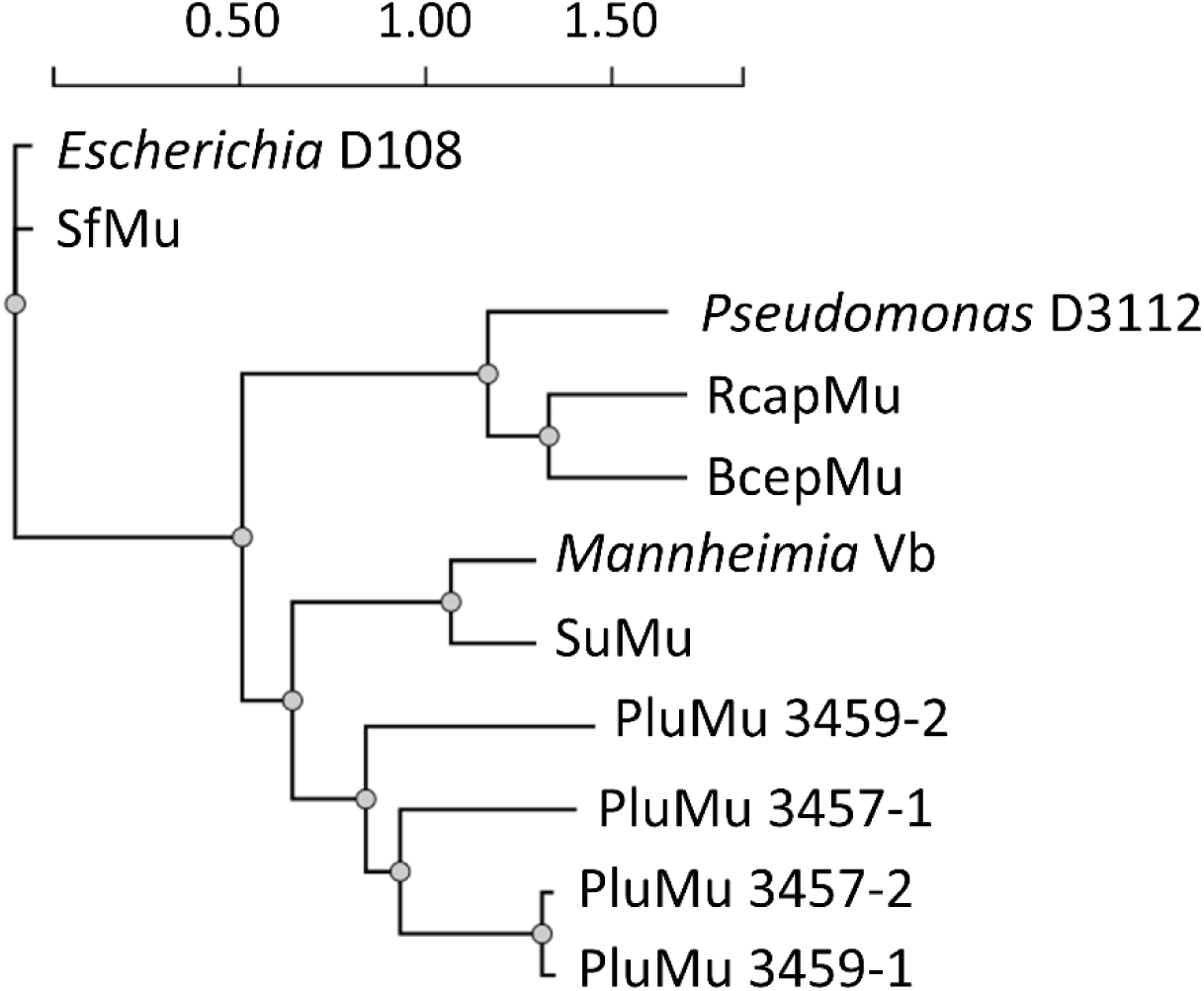
Phylogenetic tree with distances to determine the evolutionary distances between 12 Mu-like phages. All PluMu phages (especially, PluMu 4357-2 and PluMu 4359-1) are closely related. All PluMus are also closely related to phages identified in other *Pasteurellaceae*, e.g.*, Mannheimia* Vb and *Glaesserella* SuMu.

### 3.2. Molecular analyses indicate that 3457 and 3459 produce active phage

Genome analysis established the presence of Mu-like prophages in the genomes of 3457 and 3459. Initially, we used a molecular biology approach to screen for the production of PluMu. Three pairs of PCR primers were designed: two targeting PluMu genes and one a non-PluMu gene in the 3457 and 3459 genomes. Specifically, the PluMu genes targeted were those encoding the late transcriptional activator protein C which is important for the expression of morphogenic structures (Margolin et al., 1989), and encoding a tail protein. An *A. pleuropneumoniae sodC-asd* (hereafter referred to as *sodC*) amplicon [24] was used as a control to indicate the presence of bacterial gDNA. Since no amplification of *sodC* occurred in the presence of *tail* and *c* gene primers, *sodC* amplifications were done separately on the same samples. The rationale was that the DNA within intact, active virions would be protected against DNase treatment, whereas any contaminating bacterial genomic DNA (detectable by *sodC* amplification) would be digested, so amplification of *c* and *tail* genes, but not *sodC*, from DNase-treated PEG-precipitated samples would indicate the presence of active virions. PCR was performed on sample preparations of 3457, 3459, and 2331: on isolated gDNA (Figure 4A); and the PEG-precipitated filtered supernatants grown in BHI-NAD (Figure 4B). Initial PCRs established that the PluMu *c* (264 bp) and *tail* (448 bp) genes were amplified when gDNA from 3457 and 3459, but not 2331 (Figure 4A LH panel), was used as a template (Figure 4A), whereas *sodC* was amplified from gDNA of all three isolates (Figure 4A RH panel). In all cases, no amplicons were found after DNase treatment.

**Figure 4.**
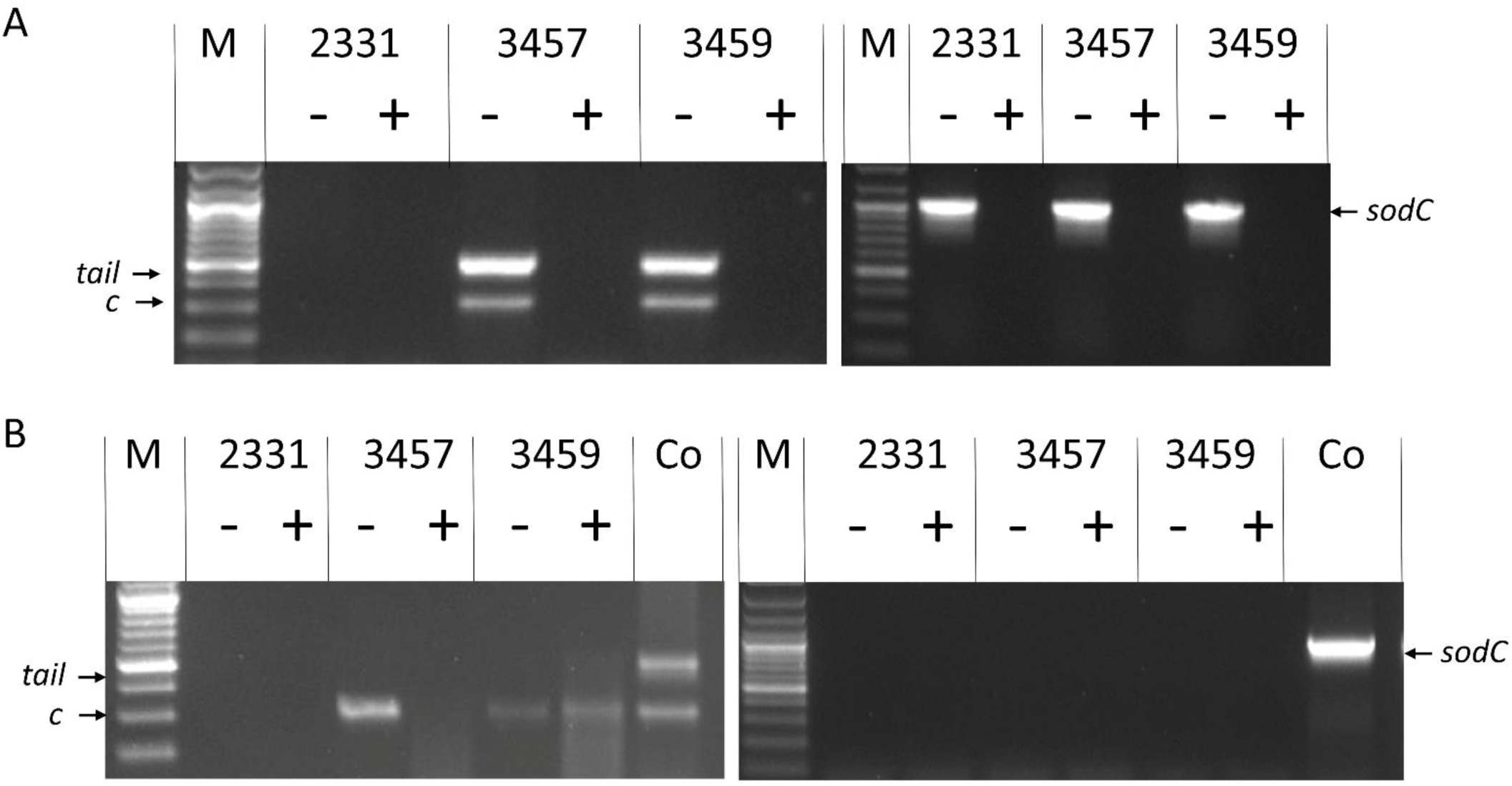
Molecular identification of phage in 3457 and 3459 in PEG-precipitated supernatants after growth in BHI-NAD. (A) Amplification of the *c* and *tail* genes present in PluMu 3457 and PluMu 3459 genomes (LH panel) and bacterial *sodC* (RH panel), using gDNA as a template without (-) and with (+) prior DNase treatment. (B) PCRs designed to amplify *c* and/or *tail* genes (LH panel) and *sodC* (RH panel) on PEG-precipitated supernatants were used as templates without (-) and with (+) prior DNase treatment. Co = 3459 gDNA. M = GeneRuler 100 bp Plus DNA Ladder (Thermo Fisher Scientific, UK).

To varying degrees, there is amplification of the *c* and *tail* genes from the PEG-precipitated samples from 3457 and 3459, both before AND after DNase treatment ((Figure 4B LH panel). indicating the presence of intact virions protecting the DNA, but no *sodC* amplification (Figure 4B RH panel) either before or after DNase treatment, indicating no contaminating gDNA was present in the samples, even before DNase treatment.

The presence of *c* and *tail*, and absence of *sodC*, amplicons in DNase-treated PEG-precipitated supernatants suggested the presence of phage particles. We also evaluated the growth of all three isolates in Grace’s Insect Medium-NAD, the rationale being that it was both defined and clear, the latter being potentially useful for observing phage by TEM. All three strains grew in NAD-supplemented Grace’s Insect Medium, although not to the same extent as in BHI-NAD broth (Figure 5). The presence of active phage particles in 3457 and 3459 was again indicated by the same *c*, *tail* (Figure 6 LH panel) and *sodC* (Fig 6. RH panel) PCR amplification results as obtained with BHI-NAD broth-grown bacteria.

**Figure 5.**
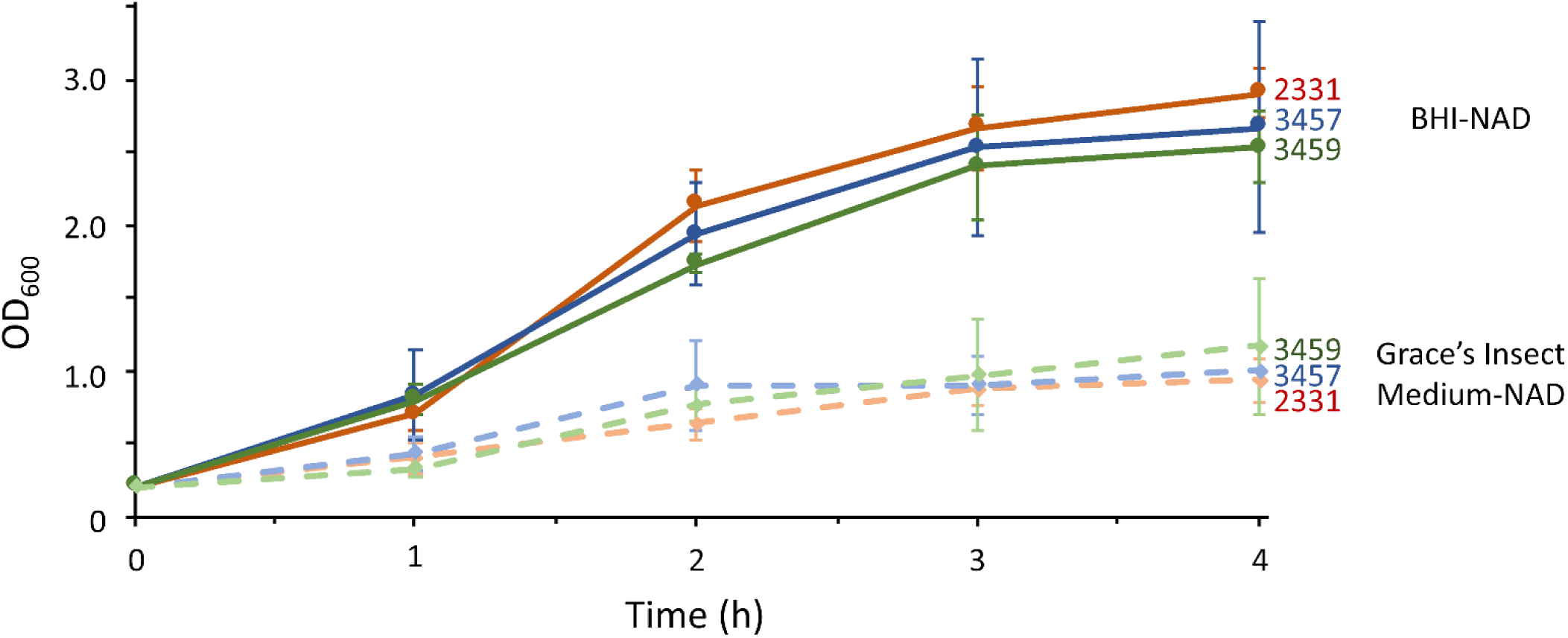
2331, 3457 and 3459 grow faster in BHI-NAD and Grace’s Insect Medium-NAD as measured by OD_600_. Error bars are mean + SD of triplicate cultures.

**Figure 6.**
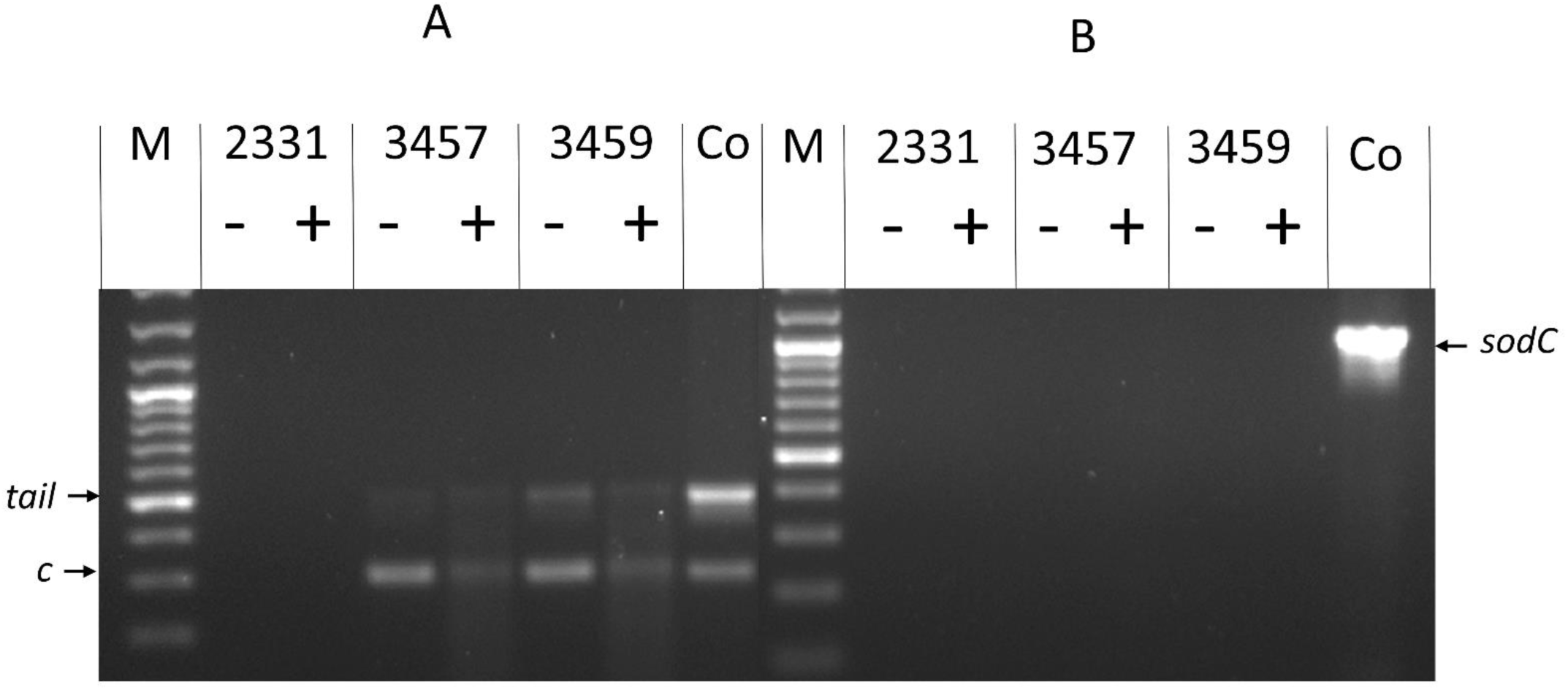
Molecular identification of active phage in PEG-precipitated supernatants of 3457 and 3459 grown in Grace’s Insect Medium-NAD. (A) Amplification of *c* and *tail* genes when PEG-precipitated supernatants were used as the template without (-) and with (+) prior DNase treatment. (B) Amplification of *A. pleuropneumoniae sodC* when PEG-precipitated supernatants were used as templates without (-) and with (+) prior DNase treatment. Co = control = 3459 gDNA. M = GeneRuler 100 bp Plus DNA Ladder (Thermo Fisher Scientific, UK).

### 3.3. Plaque formation in 3459 grown on a BHI-NAD plate

We also noticed plaque formation of 3459, but not 3457, grown on BHI-NAD agar plates (Figure 7), indicating the presence of active phage. No such plaque formation was found with 2331. Indeed, we have not observed plaque formation of any *A. pleuropneumoniae* isolate in our collection (N = c. 600) on BHI-NAD or any other agar plates.

**Figure 7.**
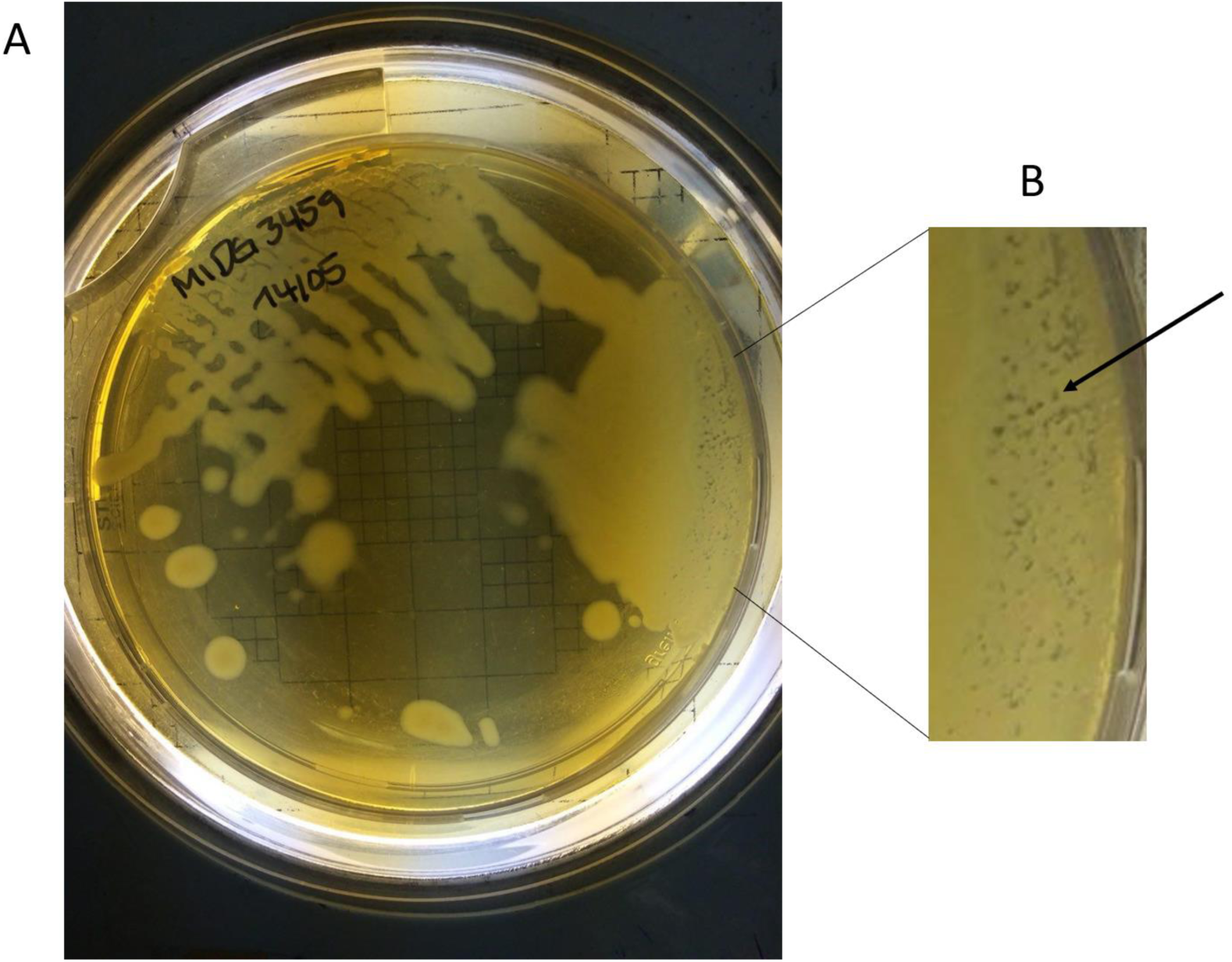
3459 grown on BHI-NAD agar plate (A) with plaque formation (B) as indicated by the arrow.

### 3.4. TEM identifies a Mu-like phage in 3459

Because of time and resource constraints and the observation of visible plaques on BHI+NAD plates, TEM analysis focused on 3459. Initial experiments analyzed samples from 3459 grown in BHI-NAD broth as the molecular analyses indicated the presence of phage (Section 3.2). Intact particles were observed but there was a high background (Figure 8A). Thus 3459 was subsequently grown in Grace’s Insect Medium-NAD, as molecular analyses had also indicated the presence of intact phage (Section 3.2). Mu-like phage particles were found visible in PEG-precipitated supernatants (Figure 8A-D). The virion enlarged in Figure 8D had a head diameter of 62.7 nm, a tail length of 202 nm, and a tail diameter of 9.4 nm The phage particles had classic Mu-like phage tail-like structures attached to icosahedral-like heads. No phage particles were found with similarly derived samples from 2331 grown in either medium.

**Figure 8.**
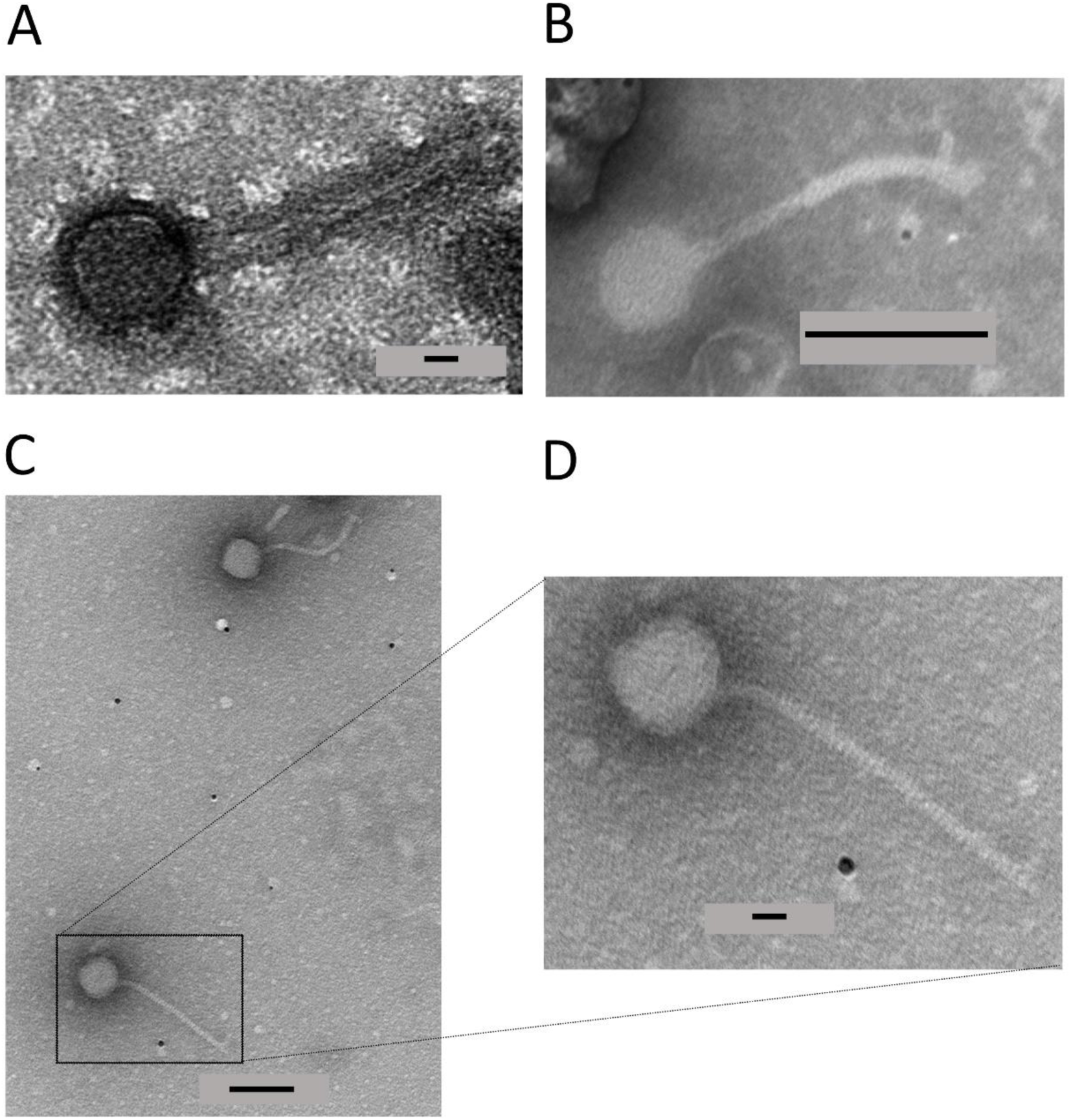
Presence of phage virions in PEG-precipitated supernatants of 3459 grown in BHI-NAD (A) or Grace’s Insect Medium-NAD (C-D). D is an enlargement of the indicated area of C. The black bars indicate distance (A = 10 nm, B = 100 nm, C = 100 nm, D = 20 nm).

## 4. Discussion

The starting point for this study was preliminary manual annotation indicating the presence of Mu-like phage genomes in two Cypriot isolates of *A. pleuropneumoniae*, 3457 and 3459 [11]. Here, we have updated that molecular analysis and provided further evidence that active Mu-like phages, which we have named PluMu, are present in those isolates. In the original manual annotation [11], only one Mu-like phage was identified in each of 3457 and 3459. Updated Phastest analysis, however, identified two Mu-like phages in each isolate. Each phage identified was ranked as “intact”, indicative of producing active phage. While their identity at the nucleotide level was poor, high levels of identity were seen at the protein level (Figure 1), especially at the 5’ ends, as illustrated by analysis of PluMu 3457-1 and 3457-2 (Figure 1). Variation in the 3’ region in Mu-like phages is a common feature, as the encoded proteins are involved in the more phage-specific functions, including tail formation and attachment. One of the genes that showed similarity to known Mu-phage (by both blastn and tblastn comparisons) was that encoding the portal protein gp29 homolog (PluMu 3457-1=CDS 31, PluMu 3457-2=CDS 19; PluMu 3459-1=CDS 19; PluMu 3459-2=CDS 11). The presence of this gene has been reported as a potential marker of virulent isolates and reference strains in *G. parasuis* [28], the bacterium where SuMu was described [14]. Further analysis will be required to determine if the PluMu gp29 homolog (or any other PluMu gene) is similarly a marker of virulent isolates. Phastest did not predict an attachment site for PluMu 3457 or PluMu 3459, unsurprising given the lack of knowledge of *A. pleuropneumoniae* phage biology. Others have used Phaster [10], or another phage identification program Prophage Hunter [9] to search *A. pleuropneumoniae* genomes for complete and/or active phage. Whole genome analysis identified 13 “complete” [10], and 21 “active” phages with 18 being most closely related to *Mannheimia* Vb [9]. In the latter study, isolate origins included Australia, Asia, Europe, and North and South America, and comprised a range of serovars, i.e., 1-2, 4-8, and 10-11. All the serovar 11 (like 3457) isolates predicted to have active phage were from Europe, although the serovar 1 (like 3459) isolate was from China [9]. Whilst neither of these studies has shown active phage, our results suggest that the presence of Mu-like phage, especially ones related to *Mannheimi*a Vb, is comparatively common in *A. pleuropneumoniae* genomes.

To determine whether 3457 or 3459 harboured active phages, as the original manual annotation and Phastest and other genomic analyses had indicated, we initially used a molecular biology approach. This involved PCR amplification of two PluMu genes (*c* and *tail*) and *A. pleuropneumoniae sodC-asd* sequences as markers of PluMu or bacterial gDNA, respectively. Despite the high specificity in PCR amplification, multiplexing all three primer pairs was unsuccessful, despite balancing melting temperatures, and checking for potential cross-dimers and sufficient size differences in amplicon size. There was some preferential amplification of the *c* gene in samples grown in Grace’s Insect Medium-NAD compared to BHI-NAD, but overall PCR analyses were reproducible across a range of different sample preparations. There was the amplification of *c* and *tail* genes in 3457 and 3459 PEG-precipitated supernatants derived from BHI-NAD and Grace’s Insect Medium-NAD media even after DNase treatment. There was no amplification of *sodC* from these samples, indicating they were free of bacterial gDNA. The results strongly suggested the presence of active PluMu, with their virion-enclosed genomes being protected from DNase treatment. To our knowledge, this is the first description of the use of defined Grace’s Insect Medium-NAD to grow *A. pleuropneumoniae*. Many groups have used Herriott’s defined medium [29] to grow *A. pleuropneumoniae* or derivatives, e.g., where the amino acid stock solution adapted from the *Neisseria* defined medium [30–32] was substituted for Herriott’s amino acid stock solution [33]. These media are comparatively more complex to make than Grace’s Insect Medium-NAD. Whilst the growth of *A. pleuropneumoniae* was not as good as in conventional BHI-NAD broth (Figure 5), further work to increase the growth rate and/or yield of cultures grown in Grace’s Insect Medium may be valuable, given the advantages of defined media in identifying gene and protein responses to changing environments.

In this study, the use of Grace’s Insect Medium was invaluable in the TEM analysis. PluMu 3459 virions were observed in PEG-precipitated supernatants derived from BHI-NAD-grown bacteria, but the background was high (Figure 8). PluMu particles were visible in PEG-precipitated supernatants derived from 3459 grown in Grace’s Insect Medium (Figure 8B-D). The isohedral heads and long tails are classic for Mu-like phage. There is some variation in the head diameter and tail length, but those identified for PluMu 3459 are comparable to those reported for other Mu-like phages. For example, the head diameter measurement of ∼62 nm is similar to bacteriophage Mu (54 nm). However, its tail length of ∼200 nm is significantly longer than that of Mu (100 nm), but comparable to BcepMu (220 nm) [34] and RcapMu (>200 nm) [35]. We are unable to categorically say whether both (or just one of) PluMu 3459-1 and PluMu 3459-2 are active phages. This question could be answered by mutation of key genes, e.g., those involved in head and/or tail formation. However, 3459, a particularly mucoid isolate, was refractory to mutant construction via conventional electroporation or natural transformation, methods we have used widely or developed [24,25,36,37]. In summary, the combination of genome and molecular analyses is consistent with the PluMu phage in 3457 and 3459 being active; confirmed by TEM for 3459.

There has been a resurgence of interest in the use of phage therapy as a strategy to treat bacterial infections, especially those caused by antimicrobial-resistant (AMR) bacteria [38]. *A. pleuropneumoniae* AMR is an increasing problem [6]. Additionally, eliminating *A. pleuropneumoniae* from herds is notoriously difficult, where complete depopulation and repopulation of a herd and deep cleansing may be required [1]. It is particularly difficult to eliminate *A. pleuropneumoniae* that are colonizing porcine tonsillar crypts, and carrier animals can be the source of acute infections in naive herds. The recognition of phage in *A. pleuropneumoniae* raises the possibility of their use to treat infection caused by AMR isolates and for eliminating tonsillar carriage. However, Mu-like phages, like PluMu, are temperate; lytic phages are used for therapy, and none have been described in *A. pleuropneumoniae*. With 19 serovars of *A. pleuropneumoniae*, it is also likely that a cocktail of phages would be required for therapy. Nonetheless, the identification of PluMu in different serovars and countries, as in this and other studies [9,10], indicates that active *A. pleuropneumoniae* lytic phage may be found in future studies. Alternative approaches such as the use of constitutively lytic phage-plasmids for therapeutic use may also be possible in the future [39].

## Supplementary materials

Phasest outputs showing CDS position, top blastp hit and E-values for PluMu phage: Table S1, PluMu 3457-1; Table S2, PluMu 3457-2; Table S3; PluMu 3459-1; Table S4 PluMu 3459-2.

## Author contributions

The following statements should be used “Conceptualization, MS, AR, JB, PL methodology, RFC, JB, YF, MS, WC, DJE, YL, JB, PL; validation, RFC, LB, YF, JB; formal analysis, RFC, LB, YF, YL, JB; investigation, RFC, LB, YF, JB; resources, MS, AR, PL; data curation, RFC, JB, YF, JB, PL; writing—original draft preparation, PL; writing—review and editing, All; visualization, All; supervision, MS, AR, JB, YL, PL; project administration, PL; funding acquisition, MS, AR, PL. All authors have read and agreed to the published version of the manuscript.

## Data availability statement

The data supporting the results and the MSc thesis of RFC is freely available from the corresponding author.

