## Supplemental Tables 1 to 4 for "PluMu - a Mu-like bacteriophage infecting *Actinobacillus pleuropneumoniae*"

### Supplementary Tables

**Table S1.** Phasest output for PluMu 3457-1 showing CDS position, top BLASTP hit and E-value.

**Region 1, total 41 CDS**

| # | CDS Position | BLAST Hit | E-Value |
|---|---|---|---|
| 1 | 66..1106 | PP_00001;integrase;phage;-;PHAGE_Mannhe_vB_MhS_1152AP2_NC_028956 | 0.0 |
| 2 | 1116..1274 | PP_00002;hypothetical protein;phage;-;PHAGE_Mannhe_vB_MhS_1152AP2_NC_028956 | 8.28e-12 |
| 3 | 1364..1585 | PP_00003;hypothetical protein;phage;-;PHAGE_Haemop_HP1_NC_001697 | 8.71e-18 |
| 4 | 1730..2230 | PP_00005;methyltransferase;phage;-;PHAGE_Mannhe_vB_MhS_535AP2_NC_028853 | 4.61e-78 |
| 5 | 2241..2960 | PP_00006;DNA methyltransferase;phage;-;PHAGE_Strept_9873_NC_047763 | 1.17e-91 |
| 6 | 3617..4414 | PP_00008;DUF2303 protein;phage;-;PHAGE_Burkho_BcepMigl_NC_019917 | 2.89e-32 |
| 7 | 4460..4819 | PP_00009;hypothetical protein;phage;-;PHAGE_Burkho_BcepMigl_NC_019917 | 6.59e-14 |
| 8 | 4882..5196 | PP_00010;hypothetical protein;phage;-;PHAGE_Burkho_BcepC6B_NC_005887 | 6.1e-10 |
| 9 | 7236..7463 | PP_00014;hypothetical protein;phage;-;PHAGE_Mannhe_vB_MhS_1152AP2_NC_028956 | 4.51e-30 |
| 10 | 7728..8003 | PP_00015;plasmid maintenance system killer HigB;phage;-;PHAGE_Mannhe_vB_MhS_535AP2_NC_028853 | 7.24e-61 |
| 11 | 8013..8303 | PP_00016;plasmid maintenance system killer HigA;phage;-;PHAGE_Mannhe_vB_MhS_535AP2_NC_028853 | 5.14e-55 |
| 12 | 8312..8479 | PP_00017;gp6, putative addiction module antidote protein;phage;-;PHAGE_Burkho_phiE12_2_NC_009236 | 1.6e-07 |
| 13 | 8671..9486 | PP_00018;putative DNA adenine methylase;phage;-;PHAGE_Geobac_E3_NC_029073 | 1.06e-20 |
| 14 | 11209..11883 | PP_00020;CI repressor;phage;-;PHAGE_Mannhe_vB_MhS_535AP2_NC_028853 | 4.72e-49 |
| 15 | 11992..12207 | PP_00021;phage protein;phage;-;PHAGE_Aggreg_S1249_NC_013597 | 5.76e-20 |
| 16 | 12266..12961 | PP_00022;uncharacterized phage-encoded protein;phage;-;PHAGE_Aggreg_S1249_NC_013597 | 1.36e-59 |
| 17 | 12958..13326 | PP_00023;hypothetical protein;phage;-;PHAGE_Mannhe_vB_MhS_587AP2_NC_028743 | 1.64e-35 |
| 18 | 13265..13981 | PP_00024;hypothetical protein;phage;-;PHAGE_Entero_mEp390_NC_019721 | 4.29e-16 |
| 19 | 13984..14508 | PP_00025;DNA N-6-adenine-methyltransferase;phage;-;PHAGE_Mannhe_vB_MhS_587AP2_NC_028743 | 1.44e-36 |
| 20 | 14703..15725 | PP_00026;hypothetical protein;phage;-;PHAGE_Shigel_SfII_NC_021857 | 5.4e-66 |
| 21 | 15797..16162 | PP_00028;endodeoxyribonuclease RusA;phage;-;PHAGE_Mannhe_vB_MhS_587AP2_NC_028743 | 1.9e-34 |
| 22 | 16152..16511 | PP_00029;antitermination protein Q;phage;-;PHAGE_Mannhe_vB_MhS_587AP2_NC_028743 | 2.73e-13 |
| 23 | 18069..19217 | PP_00032;hypothetical protein;phage;-;PHAGE_Coryne_Juicebox_NC_048070 | 7.54e-34 |
| 24 | 19541..20215 | PP_00033;antirepressor protein Ant;phage;-;PHAGE_Mannhe_vB_MhS_1152AP2_NC_028956 | 4.2e-23 |
| 25 | 21067..21426 | PP_00036;holin;phage;-;PHAGE_Mannhe_vB_MhS_587AP2_NC_028743 | 6.26e-49 |
| 26 | 21416..21862 | PP_00037;lysozyme;phage;-;PHAGE_Erwini_vB_EhrS_59_NC_048198 | 5.3e-53 |

| # | CDS Position | BLAST Hit | E-Value |
|---|---|---|---|
| 27 | 21855..22217 | PP_00038;lytic protein Rz;phage;-;PHAGE_Mannhe_vB_MhS_535AP2_NC_028853 | 6.88e-55 |
| 28 | 22162..22395 | PP_00039;lytic protein Rz1;phage;-;PHAGE_Mannhe_vB_MhS_587AP2_NC_028743 | 1.49e-31 |
| 29 | 22675..23154 | PP_00040;terminase small subunit;phage;-;PHAGE_Entero_c_1_NC_019706 | 8.02e-60 |
| 30 | 23154..25271 | PP_00041;bacteriophage DNA packaging protein;terminase, large subunit;phage;-;PHAGE_Phage_Gifsy_2_NC_010393 | 0.0 |
| 31 | 25492..27000 | PP_00043;bacteriophage portal protein;phage;-;PHAGE_Phage_Gifsy_2_NC_010393 | 0.0 |
| 32 | 27032..28990 | PP_00044;head maturation protease;phage;-;PHAGE_Entero_c_1_NC_019706 | 0.0 |
| 33 | 29064..29387 | PP_00045;hypothetical protein;phage;-;PHAGE_Entero_mEp460_NC_019716 | 1.37e-21 |
| 34 | 29685..30212 | PP_00047;bacteriophage head-tail assembly protein;Lambda gpZ homolog;phage;-;PHAGE_Phage_Gifsy_1_NC_010392 | 1.07e-27 |
| 35 | 30209..30496 | PP_00048;putative minor tail protein U;phage;-;PROPHAGE_Escher_Sakai | 6.69e-10 |
| 36 | 31415..35014 | PP_00051;putative tail component protein;phage;-;PHAGE_Pseudo_MP22_NC_009818 | 1.22e-42 |
| 37 | 35014..35754 | PP_00052;tail protein;phage;-;PHAGE_Entero_CAjan_NC_028776 | 2.27e-08 |
| 38 | 35765..36319 | PP_00053;hypothetical protein;phage;-;PHAGE_Acinet_vB_AbaS_Loki_NC_042137 | 8.55e-26 |
| 39 | 36319..36726 | PP_00054;hypothetical protein;phage;-;PHAGE_Yersin_PY54_NC_005069 | 2.53e-10 |
| 40 | 36728..38770 | PP_00055;putative tail fibre protein;phage;-;PHAGE_Acinet_vB_AbaS_Loki_NC_042137 | 3.19e-77 |
| 41 | 38815..40785 | PP_00056;tail fiber protein;phage;-;PHAGE_Klebsi_TSK1_NC_048126 | 2.95e-10 |

**Table S2**. Phasest output for PluMu 3457-2 showing CDS position, top BLASTP hit and E-value.

| # | CDS Position | BLAST Hit | E-Value |
|---|---|---|---|
| 1 | 1..423 | PP_00001;transposase;phage;-;PHAGE_Mannhe_vB_MhM_3927AP2_NC_028766 | 5.81e-15 |
| 2 | 635..847 | PP_00002;transcriptional regulatory protein;phage;-;PHAGE_Mannhe_vB_MhM_3927AP2_NC_028766 | 4.92e-28 |
| 3 | 859..2823 | PP_00003;transposase;phage;-;PHAGE_Vibrio_12B12_NC_021070 | 0.0 |
| 4 | 2883..3800 | PP_00004;transposase B;phage;-;PHAGE_Bacill_BalMu_1_NC_030945 | 3.32e-30 |
| 5 | 3804..4115 | PP_00005;hypothetical protein;phage;-;PHAGE_Mannhe_vB_MhM_3927AP2_NC_028766 | 2.03e-09 |
| 6 | 4391..4921 | PP_00007;host-nuclease inhibitor protein;phage;-;PHAGE_Haemop_SuMu_NC_019455 | 9.97e-64 |
| 7 | 5296..5484 | PP_00009;hypothetical protein;phage;-;PHAGE_Mannhe_vB_MhM_3927AP2_NC_028766 | 1.66e-08 |
| 8 | 5494..5667 | PP_00010;hypothetical protein;phage;-;PHAGE_Mannhe_vB_MhM_3927AP2_NC_028766 | 8.56e-11 |
| 9 | 6074..6691 | PP_00011;antirepressor;phage;-;PHAGE_Mannhe_vB_MhS_587AP2_NC_028743 | 1.49e-27 |
| 10 | 7070..7615 | PP_00013;hypothetical protein;phage;-;PHAGE_Vibrio_12B12_NC_021070 | 5.07e-20 |
| 11 | 7602..8156 | PP_00014;hypothetical protein;phage;-;PHAGE_Mannhe_vB_MhM_3927AP2_NC_028766 | 8.76e-55 |

| # | CDS Position | BLAST Hit | E-Value |
|---|---|---|---|
| 12 | 8301..8723 | PP_00015;putative transcription regulator;phage;-;PHAGE_Entero_Mu_NC_000929 | 3.33e-34 |
| 13 | 8812..9357 | PP_00016;putative N-acetylmuramoyl-L-alanine amidase;phage;-;PHAGE_Haemop_SuMu_NC_019455 | 6.94e-105 |
| 14 | 9984..10211 | PP_00018;DksA-like zinc finger domain containing protein;phage;-;PHAGE_Yersin_L_413C_NC_004745 | 1.05e-07 |
| 15 | 10208..10537 | PP_00019;hypothetical protein;phage;-;PHAGE_Escher_D108_NC_013594 | 1.66e-06 |
| 16 | 10543..10836 | PP_00020;hypothetical protein;phage;-;PHAGE_Escher_D108_NC_013594 | 9.11e-36 |
| 17 | 10870..11442 | PP_00021;hypothetical protein;phage;-;PHAGE_Vibrio_12B12_NC_021070 | 3.32e-56 |
| 18 | 11439..13019 | PP_00022;portal protein;phage;-;PHAGE_Vibrio_12B12_NC_021070 | 1.28e-178 |
| 19 | 13021..14586 | PP_00023;portal protein;phage;-;PHAGE_Entero_SfMu_NC_027382 | 1.15e-180 |
| 20 | 14573..15910 | PP_00024;head morphogenesis protein;phage;-;PHAGE_Entero_SfMu_NC_027382 | 4.31e-113 |
| 21 | 16115..16549 | PP_00025;putative virion morphogenesis protein;phage;-;PHAGE_Entero_Mu_NC_000929 | 1.28e-15 |
| 22 | 16780..17853 | PP_00026;putative protease protein;phage;-;PHAGE_Entero_Mu_NC_000929 | 3.95e-95 |
| 23 | 17853..18779 | PP_00027;major head protein;phage;-;PHAGE_Escher_D108_NC_013594 | 1.84e-113 |
| 24 | 19249..19656 | PP_00029;hypothetical protein;phage;-;PHAGE_Entero_SfMu_NC_027382 | 1.01e-25 |
| 25 | 19653..20078 | PP_00030;hypothetical protein;phage;-;PHAGE_Entero_SfMu_NC_027382 | 2.87e-31 |
| 26 | 20092..20835 | PP_00031;hypothetical protein;phage;-;PHAGE_Ralsto_RS138_NC_029107 | 1.05e-47 |
| 27 | 21627..21974 | PP_00033;hypothetical protein;phage;-;PHAGE_Stx2_c_Stx2a_WGPS9_NC_049923 | 6.57e-16 |
| 28 | 22030..22626 | PP_00034;hypothetical protein;phage;-;PHAGE_Stx2_c_Stx2a_WGPS9_NC_049923 | 6.08e-15 |
| 29 | 22676..23041 | PP_00035;hypothetical protein;phage;-;PHAGE_Stx2_c_Stx2a_WGPS9_NC_049923 | 2.08e-18 |

**Table S3**. Phasest output for PluMu 3459-1 showing CDS position, top BLASTP hit and E-value.

| # | CDS Position | BLAST Hit | E-Value |
|---|---|---|---|
| 1 | 1..423 | PP_00001;transposase;phage;-;PHAGE_Mannhe_vB_MhM_3927AP2_NC_028766 | 5.81e-15 |
| 2 | 635..847 | PP_00002;transcriptional regulatory protein;phage;-;PHAGE_Mannhe_vB_MhM_3927AP2_NC_028766 | 4.92e-28 |
| 3 | 859..2823 | PP_00003;transposase;phage;-;PHAGE_Vibrio_12B12_NC_021070 | 0.0 |
| 4 | 2883..3800 | PP_00004;transposase B;phage;-;PHAGE_Bacill_BalMu_1_NC_030945 | 3.32e-30 |
| 5 | 3804..4115 | PP_00005;hypothetical protein;phage;-;PHAGE_Mannhe_vB_MhM_3927AP2_NC_028766 | 2.03e-09 |
| 6 | 4391..4921 | PP_00007;host-nuclease inhibitor protein;phage;-;PHAGE_Haemop_SuMu_NC_019455 | 9.97e-64 |
| 7 | 5296..5490 | PP_00009;hypothetical protein;phage;-;PHAGE_Mannhe_vB_MhM_3927AP2_NC_028766 | 1.65e-08 |

| # | CDS Position | BLAST Hit | E-Value |
|---|---|---|---|
| 8 | 5716..6390 | PP_00011;hypothetical protein;phage;-;PHAGE_Aeromo_phiA8_29_NC_048660 | 2.71e-32 |
| 9 | 6883..7431 | PP_00014;hypothetical protein;phage;-;PHAGE_Vibrio_12B12_NC_021070 | 1.62e-20 |
| 10 | 7418..7972 | PP_00015;hypothetical protein;phage;-;PHAGE_Mannhe_vB_MhM_3927AP2_NC_028766 | 8.76e-55 |
| 11 | 8117..8539 | PP_00016;putative transcription regulator;phage;-;PHAGE_Entero_Mu_NC_000929 | 3.33e-34 |
| 12 | 8628..9173 | PP_00017;putative N-acetylmuramoyl-L-alanine amidase;phage;-;PHAGE_Haemop_SuMu_NC_019455 | 6.94e-105 |
| 13 | 9180..9404 | PP_00018;hypothetical protein;phage;-;PHAGE_Haemop_SuMu_NC_019455 | 2.73e-16 |
| 14 | 9800..10027 | PP_00020;DksA-like zinc finger domain containing protein;phage;-;PHAGE_Yersin_L_413C_NC_004745 | 1.05e-07 |
| 15 | 10027..10353 | PP_00021;hypothetical protein;phage;-;PHAGE_Escher_D108_NC_013594 | 1.62e-06 |
| 16 | 10359..10652 | PP_00022;hypothetical protein;phage;-;PHAGE_Escher_D108_NC_013594 | 9.11e-36 |
| 17 | 10686..11258 | PP_00023;hypothetical protein;phage;-;PHAGE_Vibrio_12B12_NC_021070 | 3.32e-56 |
| 18 | 11255..12835 | PP_00024;portal protein;phage;-;PHAGE_Vibrio_12B12_NC_021070 | 1.28e-178 |
| 19 | 12837..14402 | PP_00025;portal protein;phage;-;PHAGE_Entero_SfMu_NC_027382 | 1.15e-180 |
| 20 | 14389..15726 | PP_00026;head morphogenesis protein;phage;-;PHAGE_Entero_SfMu_NC_027382 | 4.31e-113 |
| 21 | 15931..16365 | PP_00027;putative virion morphogenesis protein;phage;-;PHAGE_Entero_Mu_NC_000929 | 1.28e-15 |
| 22 | 16596..17669 | PP_00028;putative protease protein;phage;-;PHAGE_Entero_Mu_NC_000929 | 3.95e-95 |
| 23 | 17669..18595 | PP_00029;major head protein;phage;-;PHAGE_Escher_D108_NC_013594 | 1.84e-113 |
| 24 | 19065..19472 | PP_00031;hypothetical protein;phage;-;PHAGE_Entero_SfMu_NC_027382 | 1.01e-25 |
| 25 | 19469..19894 | PP_00032;hypothetical protein;phage;-;PHAGE_Entero_SfMu_NC_027382 | 2.87e-31 |
| 26 | 19908..20651 | PP_00033;hypothetical protein;phage;-;PHAGE_Ralsto_RS138_NC_029107 | 1.05e-47 |
| 27 | 20709..21119 | PP_00034;hypothetical protein;phage;-;PHAGE_Pseudo_JBD93_NC_030918 | 7.75e-22 |
| 28 | 21565..24924 | PP_00036;tail length tape measure protein;phage;-;PHAGE_Entero_mEp390_NC_019721 | 3.74e-40 |
| 29 | 24924..25922 | PP_00037;hypothetical protein;phage;-;PHAGE_Pseudo_JBD67_NC_042135 | 8.49e-42 |
| 30 | 25925..26890 | PP_00038;hypothetical protein;phage;-;PHAGE_Pseudo_vB_PaeS_PM105_NC_028667 | 2.3e-40 |
| 31 | 26893..28575 | PP_00039;hypothetical protein;phage;-;PHAGE_Pseudo_DMS3_NC_008717 | 2.52e-108 |
| 32 | 28612..29421 | PP_00040;BR0599 family protein;phage;-;PHAGE_Salmon_KFS_SE1_NC_048683 | 3.29e-56 |
| 33 | 29439..29690 | PP_00041;conserved tail assembly protein;phage;-;PHAGE_Burkho_BcepNazgul_NC_005091 | 1.08e-19 |
| 34 | 29695..29898 | PP_00042;tail assembly protein;phage;-;PHAGE_Vibrio_VpKK5_NC_026610 | 1.73e-16 |
| 35 | 29898..32681 | PP_00043;conserved tail assembly protein;phage;-;PHAGE_Burkho_BcepNazgul_NC_005091 | 6.5e-142 |
| 36 | 33040..33435 | PP_00044;phage protein;phage;-;PHAGE_Strept_phiBHN167_NC_022791 | 5.03e-47 |
| 37 | 33413..33796 | PP_00045;putative C-5 cytosine-specific DNA methylase;phage;-;PHAGE_Strept_phiNJ2_NC_019418 | 1.7e-38 |
| 38 | 33789..34625 | PP_00046;hypothetical protein;phage;-;PHAGE_Haemop_SuMu_NC_019455 | 2.04e-137 |

| # | CDS Position | BLAST Hit | E-Value |
|---|---|---|---|
| 39 | 35450..36028 | PP_00048;DNA-directed RNA polymerase specialized sigma subunit;phage;-;PHAGE_Sinorh_phiM9_NC_028676 | 2.77e-08 |

**Table S4**. Phasest output for PluMu 3459-2 showing CDS position, top BLASTP hit and E-value.

| # | CDS Position | BLAST Hit | E-Value |
|---|---|---|---|
| 1 | 9118..11088 | PP_00011;tail fiber protein;phage;-;PHAGE_Klebsi_TSK1_NC_048126 | 2.95e-10 |
| 2 | 11133..13175 | PP_00012;putative tail fibre protein;phage;-;PHAGE_Acinet_vB_AbaS_Loki_NC_042137 | 3.19e-77 |
| 3 | 13177..13584 | PP_00013;hypothetical protein;phage;-;PHAGE_Yersin_PY54_NC_005069 | 2.53e-10 |
| 4 | 13584..14138 | PP_00014;hypothetical protein;phage;-;PHAGE_Acinet_vB_AbaS_Loki_NC_042137 | 8.55e-26 |
| 5 | 14138..14878 | PP_00015;tail protein;phage;-;PHAGE_Entero_CAjan_NC_028776 | 1.43e-13 |
| 6 | 14878..18477 | PP_00016;putative tail component protein;phage;-;PHAGE_Pseudo_MP22_NC_009818 | 1.22e-42 |
| 7 | 19260..19676 | PP_00019;putative minor tail protein U;phage;-;PROPHAGE_Escher_Sakai | 3.79e-14 |
| 8 | 19673..20200 | PP_00020;bacteriophage head-tail assembly protein;Lambda gpZ homolog;phage;-;PHAGE_Phage_Gifsy_1_NC_010392 | 1.07e-27 |
| 9 | 20498..20821 | PP_00022;hypothetical protein;phage;-;PHAGE_Entero_mEp460_NC_019716 | 1.37e-21 |
| 10 | 20895..22853 | PP_00023;head maturation protease;phage;-;PHAGE_Entero_c_1_NC_019706 | 0.0 |
| 11 | 22885..24393 | PP_00024;bacteriophage portal protein;phage;-;PHAGE_Phage_Gifsy_2_NC_010393 | 0.0 |
| 12 | 24614..26731 | PP_00026;bacteriophage DNA packaging protein;terminase, large subunit;phage;-;PHAGE_Phage_Gifsy_2_NC_010393 | 0.0 |
| 13 | 26731..27210 | PP_00027;terminase small subunit;phage;-;PHAGE_Entero_c_1_NC_019706 | 8.02e-60 |
| 14 | 27490..27723 | PP_00028;lytic protein Rz1;phage;-;PHAGE_Mannhe_vB_MhS_587AP2_NC_028743 | 1.49e-31 |
| 15 | 27668..28030 | PP_00029;lytic protein Rz;phage;-;PHAGE_Mannhe_vB_MhS_535AP2_NC_028853 | 6.88e-55 |
| 16 | 28023..28469 | PP_00030;lysozyme;phage;-;PHAGE_Erwini_vB_EhrS_59_NC_048198 | 5.3e-53 |
| 17 | 28459..28818 | PP_00031;holin;phage;-;PHAGE_Mannhe_vB_MhS_587AP2_NC_028743 | 6.26e-49 |
| 18 | 29670..30344 | PP_00034;antirepressor protein Ant;phage;-;PHAGE_Mannhe_vB_MhS_1152AP2_NC_028956 | 4.2e-23 |
| 19 | 30668..31816 | PP_00035;hypothetical protein;phage;-;PHAGE_Coryne_Juicebox_NC_048070 | 1.02e-33 |
| 20 | 33374..33733 | PP_00038;antitermination protein Q;phage;-;PHAGE_Mannhe_vB_MhS_587AP2_NC_028743 | 2.73e-13 |
| 21 | 33723..34088 | PP_00039;endodeoxyribonuclease RusA;phage;-;PHAGE_Mannhe_vB_MhS_587AP2_NC_028743 | 1.9e-34 |
| 22 | 34160..35182 | PP_00041;hypothetical protein;phage;-;PHAGE_Shigel_SfII_NC_021857 | 5.4e-66 |
| 23 | 35175..35393 | PP_00042;hypothetical protein;phage;-;PHAGE_Mannhe_vB_MhS_535AP2_NC_028853 | 2.78e-08 |
| 24 | 35377..35901 | PP_00043;DNA N-6-adenine-methyltransferase;phage;-;PHAGE_Mannhe_vB_MhS_587AP2_NC_028743 | 1.44e-36 |
| 25 | 35904..36620 | PP_00044;hypothetical protein;phage;-;PHAGE_Entero_mEp390_NC_019721 | 4.29e-16 |
| 26 | 36559..36927 | PP_00045;hypothetical protein;phage;-;PHAGE_Mannhe_vB_MhS_587AP2_NC_028743 | 1.64e-35 |

| # | CDS Position | BLAST Hit | E-Value |
|---|---|---|---|
| 27 | 36924..37619 | PP_00046;uncharacterized phage-encoded protein;phage;-;PHAGE_Aggreg_S1249_NC_013597 | 1.36e-59 |
| 28 | 37678..37893 | PP_00047;phage protein;phage;-;PHAGE_Aggreg_S1249_NC_013597 | 5.76e-20 |
| 29 | 38002..38676 | PP_00048;CI repressor;phage;-;PHAGE_Mannhe_vB_MhS_535AP2_NC_028853 | 4.72e-49 |
| 30 | 40399..41214 | PP_00050;putative DNA adenine methylase;phage;-;PHAGE_Geobac_E3_NC_029073 | 1.06e-20 |
| 31 | 41406..41573 | PP_00051;gp6, putative addiction module antidote protein;phage;-;PHAGE_Burkho_phiE12_2_NC_009236 | 1.6e-07 |
| 32 | 41582..41872 | PP_00052;plasmid maintenance system killer HigA;phage;-;PHAGE_Mannhe_vB_MhS_535AP2_NC_028853 | 5.14e-55 |
| 33 | 41882..42157 | PP_00053;plasmid maintenance system killer HigB;phage;-;PHAGE_Mannhe_vB_MhS_535AP2_NC_028853 | 7.24e-61 |
| 34 | 42422..42649 | PP_00054;hypothetical protein;phage;-;PHAGE_Mannhe_vB_MhS_1152AP2_NC_028956 | 4.51e-30 |
| 35 | 44689..45003 | PP_00058;hypothetical protein;phage;-;PHAGE_Burkho_BcepC6B_NC_005887 | 6.1e-10 |
| 36 | 45066..45425 | PP_00059;hypothetical protein;phage;-;PHAGE_Burkho_BcepMigl_NC_019917 | 6.59e-14 |
| 37 | 45471..46268 | PP_00060;DUF2303 protein;phage;-;PHAGE_Burkho_BcepMigl_NC_019917 | 2.89e-32 |
| 38 | 46861..47664 | PP_00062;hypothetical protein;phage;-;PHAGE_Strept_phiARI0468_4_NC_031915 | 1.96e-101 |
| 39 | 47675..48175 | PP_00063;methyltransferase;phage;-;PHAGE_Mannhe_vB_MhS_535AP2_NC_028853 | 4.61e-78 |
| 40 | 48320..48541 | PP_00065;hypothetical protein;phage;-;PHAGE_Haemop_HP1_NC_001697 | 8.71e-18 |
| 41 | 48631..48789 | PP_00066;hypothetical protein;phage;-;PHAGE_Mannhe_vB_MhS_1152AP2_NC_028956 | 8.28e-12 |
| 42 | 48799..49839 | PP_00067;integrase;phage;-;PHAGE_Mannhe_vB_MhS_1152AP2_NC_028956 | 0.0 |
